# Principled simulation of agent-based models in epidemiology

**DOI:** 10.1101/2020.12.21.423765

**Authors:** Sean L. Wu, Andrew J. Dolgert, Joseph A. Lewnard, John M. Marshall, David L. Smith

## Abstract

After more than a century of sustained work by mathematicians, biologists, epidemiologists, probabilists, and other experts, dynamic models have become a vital tool for understanding and describing epidemics and disease transmission systems. Such models fulfill a variety of crucial roles including data integration, estimation of disease burden, forecasting trends, counterfactual evaluation, and parameter estimation. These models often incorporate myriad details, from age and social structure to inform population mixing patterns, commuting and migration, and immunological dynamics, among others. This complexity can be daunting, so many researchers have turned to stochastic simulation using agent-based models. Developing agent-based models, however, can present formidable technical challenges. In particular, depending on how the model updates state, unwanted or even unnoticed approximations can be introduced into a simulation model. In this article, we present computational methods for approximating continuous time discrete event stochastic processes based on a discrete time step to speed up complicated simulations which also converges to the true process as the time step goes to zero. Our stochastic models is constructed via hazard functions, and only those hazards which are dependent on the state of other agents (such as infection) are approximated, whereas hazards governing dynamics internal to an agent (such as immune response) are simulated exactly. By partitioning hazards as being either dependent or internal, a generic algorithm can be presented which is applicable to many models of contagion processes, with natural areas of extension and optimization.

**Author summary:** Stochastic simulation of epidemics is crucial to a variety of tasks in public health, encompassing intervention evaluation, trend forecasting, and estimation of epidemic parameters, among others. In many situations, due to model complexity, time constraints, unavailability or unfamiliarity with existing software, or other reasons, agent-based models are used to simulate epidemic processes. However, many simulation algorithms are *ad hoc*, which may introduce unwanted or unnoticed approximations. We present a method to build approximate, agent-based models from mathematical descriptions of stochastic epidemic processes which will improve simulation speed and converge to exact simulation techniques in limiting cases. The simplicity and generality of our method should be widely applicable to various problems in mathematical epidemiology and its connection to other methods developed in chemical physics should inspire future work and elaboration.

## Introduction

Stochastic simulation of complex mathematical models is a vital tool for understanding and describing disease transmission systems. While early efforts by probabilists Bailey and Kendall [1, 2], and the biochemist and physician-epidemiologist pair Kermack and McKendrick [3, 4] were highly mathematical in nature, they were at best, coarse approximations of natural processes. In the century hence, disease transmission models have incorporated myriad details including age structure, commuting, migration, and within-host immunology to better represent our understanding of how these systems function. Complex simulation models have been used to great effect in understanding rapidly changing epidemic situations, such as the outbreak of a novel pathogen, or events which require immediate response, such as the 2001 veterinary epidemic of hand foot and mouth disease in the UK [5]. When addressing urgent public health crises, complex models of epidemic processes can be invaluable tools, serving as platforms for data integration [6], estimation of current burden [7], forecasting trends [8], evaluation of intervention strategies and counterfactual scenarios [9], among other roles.

These models are often formulated as compartmenal models and simulated as stochastic jump processes, where a jump is a change in an individual’s epidemiological state. Sample paths (trajectories) are piecewise constant, and change only due the to occurrence of discrete events that change the epidemiological state of individuals (jumps). When jumps are only allowed to occur after exponentially distributed intervals, the process is a continuous-time Markov chain (CTMC). Because the modeler is free to design the states, state transitions, and associated distribution of inter-jump intervals as they see fit, this class of models can be directly specified from the results of survival analysis [10], allowing close alignment to empirical data and standard survival models. In general, jump processes are also valued for their deep connection to deterministic models (*e*.*g*., compartmental models) in limiting cases which can aid model verification; such results are well known for CTMC models [11, 12] and exist for some non-Markovian extensions [13]. Furthermore, these models benefit from over a half-century of rigorous mathematical and algorithmic study [14], especially in chemical kinetics, physics, and operations research communities, which has led to a plethora of publicly available algorithms to sample trajectories from such processes, as well as techniques for statistical inference and model fitting.

The complexity of these models, however, frustrates analytic approaches and can even thwart straightforward application of classic simulation techniques, making quick development and application challenging, especially in epidemic response situations. Incorporating non-Markovian dynamics is still difficult in most simulation frameworks, and because the majority of “industrial strength” simulation software is designed for relatively straightforward chemical reaction networks [15], peculiarities of epidemiological simulation such as highly nonlinear force of infection terms, age and location-based mixing, immunological dynamics, and other elaborations make direct utilization of these software difficult. While there exist some open-source software for simulation of epidemiological dynamics, many frameworks are restricted to simple compartmental models, or may have a significant enough learning curve that they are simply not an option when a model must be developed in a matter of days [16, 17]. In addition, researchers may require specific forms of model output that can be difficult for software packages to support.

Beacuse of their expressive power, agent-based models (ABM) are often the most straightforward way to turn a whiteboard description of a complex system into useable code, and are a viable alternative to other methods of representing a model. Many ABMs are developed as bespoke programs for a specific analysis, but the technical complexity of implementation means that a variety of subjective decisions may be made when writing simulation code. Not all design choices will lead to simulation algorithms that necessarily have a limiting interpretation as some stochastic jump process, valuable for both model verification and model modification based on the formal rules of the stochastic process.

Here we describe a method to construct approximate ABM representations of stochastic models in which agents with arbitrarily complex internal dynamics interact through discrete events in continuous time. Our method relies on specification of a discrete time step of size Δ*t*, over which interactions *between* agents are approximated (dependent events); dynamics *within* an agent (internal events) are still simulated exactly in continuous time. A significant contribution in this work is presenting a generic algorithm to simulate systems that are relevant to a wide class of epidemiological models, via approximation of dependent events which can help speed up even highly complex ABMs. We also demonstrate that our algorithm approaches the true continuous time jump process as Δ*t* → 0. To verify our method and provide numerical comparisons to exact stochastic simulation, we use our algorithm to simulate a Markovian and non-Markovian SIR (Susceptible-Infectious-Recovered) model. We conclude with a discussion of strengths and weaknesses of our approach, as well as fruitful next steps to generalize our method. We hope that our method gives mathematical epidemiologists considerable freedom in designing a model to fit their needs, and that by approximating dependent events, even highly complex models can be feasible to simulate. We also expect our method will be of interest to researchers in ecology, demography, and the quantitative social sciences.

## Materials and methods

### Hazard Rates in Stochastic Simulation

Formally, a jump process is a stochastic process, *X* whose trajectories (sample paths) are piecewise constant functions of time over a countable set of states, *S* so that *X*(*t*) ∈*S, t* ≥ 0. Here we restrict ourselves to considering time-homogeneous processes, so that only the dwell time in the current state is relevant for the process. In order to sample trajectories from *X*, we must specify hazard functions *λ*_*j*_(*τ, s*) associated with each event *j* ∈ {1, …, *M*}. The hazard functions tell us the conditional probability of *j* occurring in the next infinitesimal time interval [*τ, τ* + *dt*), if the process has dwelled in state *s* ∈*S* for some time *τ* (in the case where there is no dependence on *τ*, the hazard will be a constant value and *X* is a CTMC). When *j* occurs, it is allowed to change the state *X* in some way. Given a set of *M* events, simulation consists of sampling when the next event occurs, which event was it, updating state appropriately, recalculating hazards that change, and repeating this process [18]. The state space can be defined implicitly, and can be (countably) infinite, so long as only a finite number of events have non-zero hazards at any time.

To define the process that the ABM will sample from, we let *X*(*t*) = (*s*_1_, …, *s*_*n*_) be a vector where *n* is the number of agents being simulated, and each *s*_*h*_ is the state of person *h*. Expanding state space in this was to achieve an agent-based representation is known as disaggregation, and may be used to study ABMs via the technical condition of lumpability [19]. Then, any events which affect person *h* and whose hazard function depends on more elements of *X* than only *s*_*h*_ is a dependent event; events which affect person *h* and whose hazard function is only allowed to depend on *s*_*h*_ is an internal event. Consider the recovery event in an SIR model. The recovery event for person *h* does not need to know about the states of anyone else in order to calculate the hazard of recovery and so is an internal event. However, the infection event for individual *h does* require knowing the state of other agents in order to compute the hazard, and therefore is a dependent event.

To draw approximate trajectories from the ABM requires a choice of time step Δ*t*, over which hazards for each agent’s dependent events may only use information from other agents at the start of the step, ignoring changes which occur *during* the time step. Put another way, agents only exchange information at the start of each time step. Then over a time step beginning at time *t*, a susceptible agent *i* is subject to infection hazard (force of infection, hereafter FOI) *β*(*u*)*I*(*t*) for *u* ∈ [*t, t* + Δ*t*), where *β*(*u*) is the effective contact rate [20]. Note that the contact rate could be allowed to depend on time, and that the approximation is in setting the number of infected individuals to a constant value (the number at time *t*). Dependent events can be simulated via rejection sampling. Internal events, by definition will not depend on any other individuals and therefore will not be approximated. This type of simulation is reminiscent of tau-leaping techniques [21], but whereas in standard tau-leaping *all* hazard functions are approximated as constants over the time step, our method only approximates dependent events, and internal events are simulated exactly.

We postpone a complete description of our method to section Simulation Algorithm, and first demonstrate that the accept-reject sampling we use for sampling dependent event times can draw correct times in a base case.

### Approximation of Hazard Rates

Consider the a single susceptible agent who is subject to a single event, infection. Let *τ*_*S*→*I*_ be a random variable giving the time at which this agent becomes infected, *tmax* be the end of the current time step, [*tmax* − Δ*t, tmax*), and *λ* be the FOI which is valid over that time step.

In our model, each agent stores its current state *s*_*h*_, the time at which it entered that state *tnow*, and the next scheduled event time *tnext* and stat 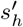. For all internal events, simulation is identical to classic discrete event simulation techniques. While the agent’s next scheduled event time is less than *tmax* the agent will update their state and time according to that scheduled event, and sample a new state and time. If the new event time is greater than *tmax*, the update does not occur until the time step in which that event falls.

In the case of infection, a dependent event, if we naively sampled the time of infection as *τ*_*S*→*I*_ ∼ *tnow* + *Exp*(*λ*), we would be ignoring future stochastic changes in FOI, which could change at *tmax*. Crucially, from the perspective of this agent, the FOI is a stochastic quantity, because it depends on the states of other agents. To develop a reasonable approximation, one needs to implement a rejection algorithm to sample *τ*_*S*→*I*_.

The sampling is described graphically in Fig 1 A. During this time step, *λ* is constant (red region), and the agent samples a *putative* time to infection 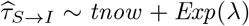. Note that there is no restriction *tnow* be equal to the start of the time step, because it could have been a randomly sampled quantity from previous events. If 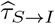 falls within the remaining time for which that hazard is valid (purple region of size *tmax* − *tnow*, which is to say 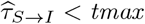), then we accept the sample and infect the agent at time 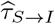. The probability of the putative time being accepted is 1 − *e*^−*λ*(*tmax* − *tnow*)^.

**Fig 1.**
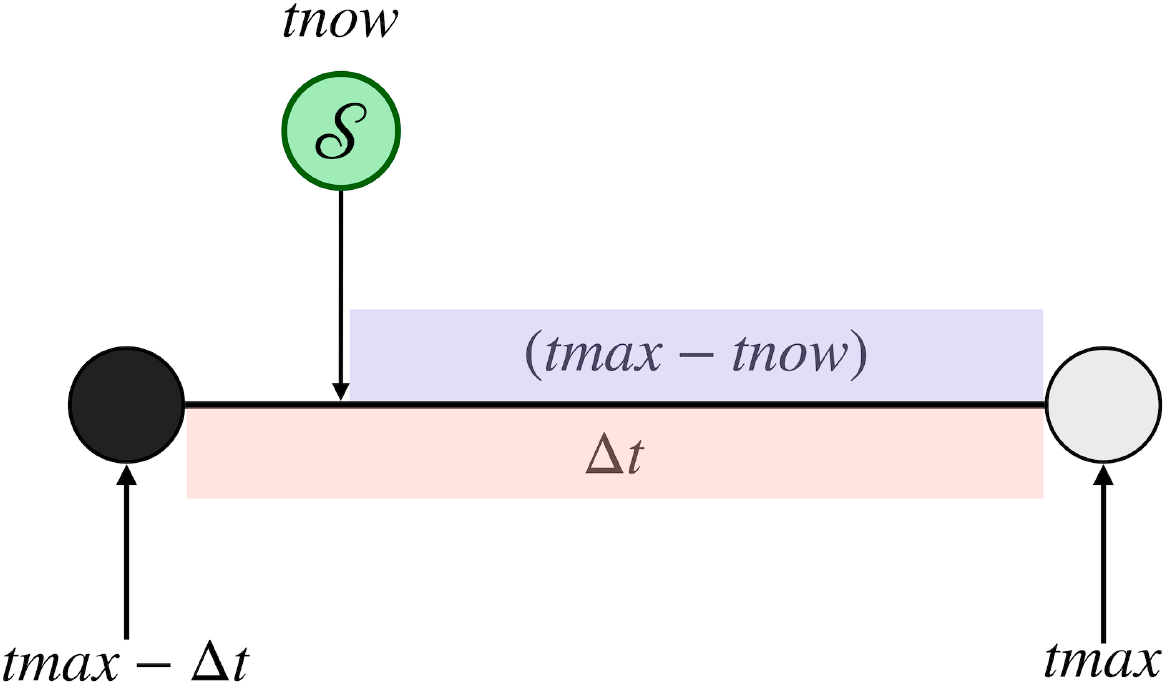
Approximate infection hazard over a time step. The left filled circle and unfilled right circle indicate that time steps are closed on the left and open on the right.

If however we reject the putative time 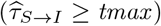 then we set the agents next event time to be *tmax*, and do not change the agent’s state. Then, on the next time step, the agent sets their current time to the start of that time step and resamples 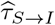 using the *newly updated* FOI *λ*′. The probability of acceptance is 1 − *e*^−*λ*′Δ*t*^.

### Accept-reject Algorithm for Piecewise Constant Hazard Rates

When Δ*t* → 0, the accept-reject algorithm for time of infection becomes exact. To demonstrate this, first consider the case where the FOI is a deterministic quantity, and, furthermore, that it is a constant. Then the probability of acceptance on any time step of size Δ*t* is *p* = 1 − *e*^−*λ*Δ*t*^. Consequently, the expected number of rejections prior to the acceptance is given as 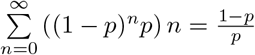. The expected number of trials including the acceptance is thus 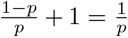, or the expectation of a Geometric random variable. Note that this is not quite the same as the expected value of the time to infection, which is 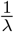. The value 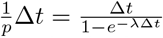”overshoots” because the acceptance could have occurred at any point in the final time step, not just at the end.

Now, conditional on being accepted during trial *n* (that is, prior to its conclusion), the time *τ* at which the acceptance occurs within the time step [(*n* − 1)Δ*t, n*Δ*t*) is given by an upper truncated Exponential distribution (S1 Fig). This random variable has expectation 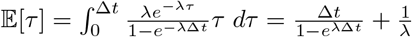. Then the time at which the event occurs should be equal to the number of rejections, each scaled by the time step size, plus this quantity, which is: 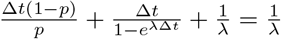, so we show that the accept-reject algorithm recovers precisely the same average waiting time as the Exponential random variate.

If instead of constant, the FOI is a deterministic piecewise constant function such that *λ*_*n*_ is the value of the FOI between [*n*Δ*t*, (*n* + 1)Δ*t*) similar reasoning applies but the number of trials needed no longer follows a Geometric distribution. The time of acceptance within the acceptance interval however, still follows a truncated Exponential, because it is conditioned on the infection event occurring in that time step of (piecewise) constant hazard. In this case, sampling *τ*_*S*→*I*_ is equivalent to sampling the first event time of an nonhomogeneous Poisson process (NHPP) with intensity function *λ*(*t*).

From [22], one way to sample from such a process is by inversion of the distribution of inter-event times, which has distribution function 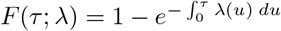. To sample the first event time, one should draw a uniform random number *u* between [0, 1) and solve so that *τ* is the first event time: 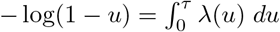. If the cumulative hazard function 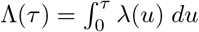 is especially easy to invert, then by the random time change (RTC) theorem, we can also sample from the distribution of *t* by sampling *v* from a unit rate Exponential distribution and then solving *τ* = Λ^−1^(*v*) [23].

When FOI is piecewise constant, Λ(*τ*) will be piecewise linear so inversion of the cumulative hazard will be the most straightforward sampling method. To compare the accept-reject algorithm with inversion sampling of NHPP first event times, we discretized a continuous intensity function 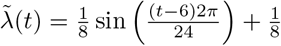. The function has a period of 24 hours, with a maximum amplitude of 0.25 (corresponding to 1/4 events an hour) with a vertical shift so that it is non-negative and a phase shift of 6 hours such that the maximum intensity occurs each day at noon and minimum intensity at midnight (S2 Fig). The integrated continuous intensity is 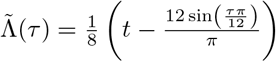.

In Fig 2 we show that the accept-reject sampler for first event times of NHPPs is equivalent to exact inversion methods given in [22, 23]. In both panels the red curve is the density function of first event times calculated directly from 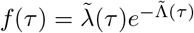, and we drew 10^6^ samples using each algorithm to construct the histograms. We see that both the accept-reject algorithm and integrated hazard sampling are sampling from the correct density function.

**Fig 2.**
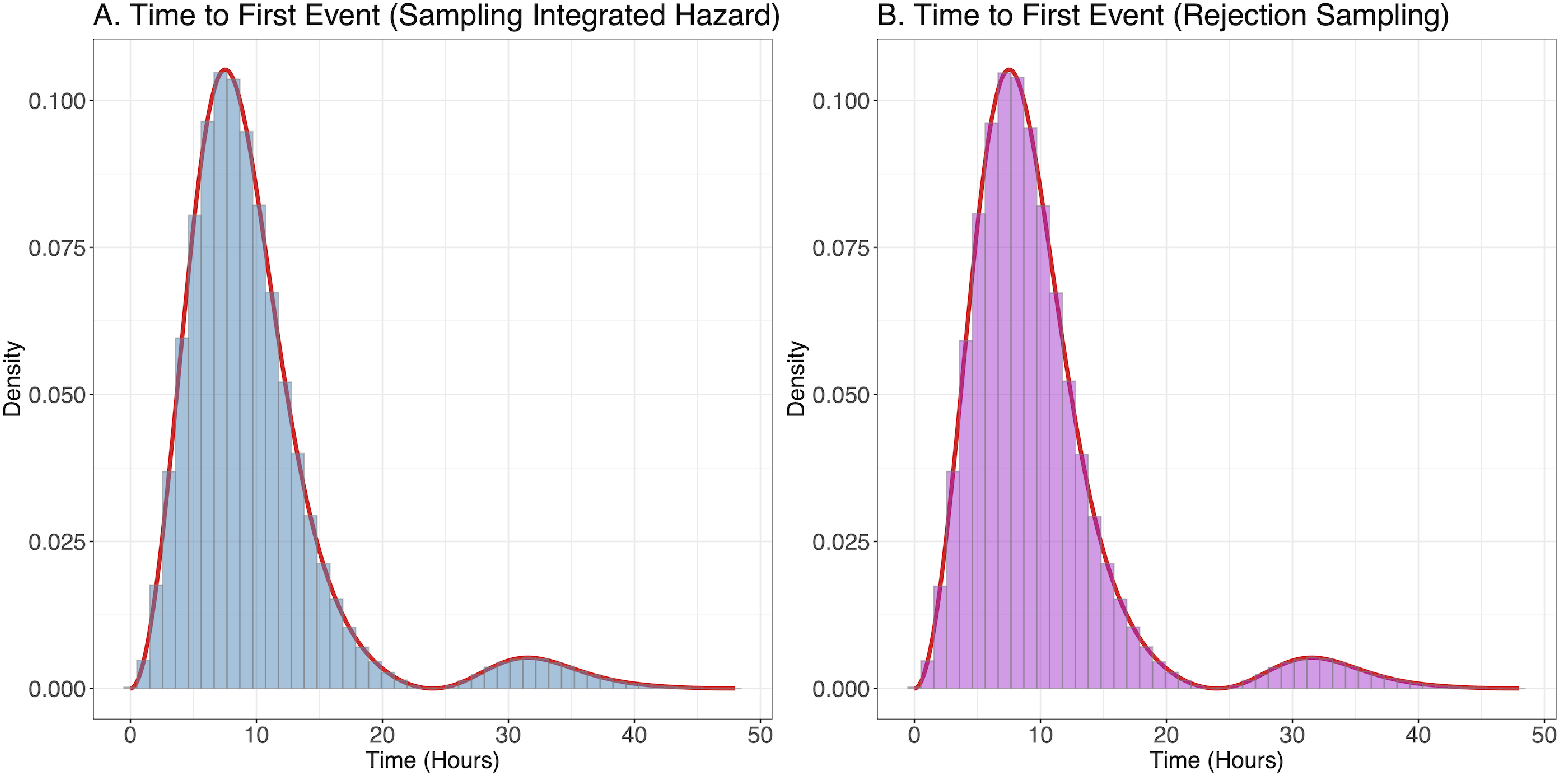
Comparison of Rejection Sampler and Direct Inversion for sampling first event times. Panel A: using integrated hazard to sample, Panel B: using accept-reject algorithm to sample. Red curves in both panels are the exact density, from numerical integration.

In a full model with interacting agents, each agent’s FOI depends on the state of other agents, and the piecewise constant approximation means that on the [*n*Δ*t*, (*n* + 1)Δ*t*) time step that FOI is computed at *n*Δ*t* and remains constant until the next time step at time (*n* + 1)Δ*t*, when agents may exchange information. Let us consider what happens as we let Δ*t* → 0. Because the model is a set of simple counting processes in continuous time, meaning that only jumps of size 1 are allowed, the probability of two events occurring simultaneously is zero [24]. So when Δ*t* is near zero, we expect that on each time step either 0 or 1 event will occur, regardless of how many agents are in the system. With infinitesimal time steps, after a single event occurs, the FOI for all agents will be updated *immediately* and the algorithm becomes exact.

### Simulation Algorithm

Our agent-based simulation algorithm for stochastic epidemic models where each agent is subject to a single dependent event, infection, is given in pseudocode below. Generalization to multiple dependent events (such as multiple strains or routes of transmission, for example) can be easily accommodated by keeping track of multiple force of infection (hazard) terms for each agent, and letting the accept-reject algorithm sample the first event time over all competing dependent events. Each agent in the population being simulated (*h* ∈ {1, …, *n*}) stores (at minimum) the following pieces of information: their current state *s*_*h*_, next state 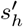, current time *tnow*_*h*_, and next time *tnext*_*h*_.

1. Initialize. For each agent *h*, set *tnow*_*h*_ = *tnext*_*h*_ = 0 and set their initial state 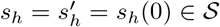. Set system time *t* = 0.
2. Set *tmax* = *t* + Δ*t*.
3. Compute the force of infection on each agent *λ*_*h*_.
4. Simulate each human’s trajectory between [*tnow*_*h*_, *tmax*):
  - While *tnext*_*h*_ < *tmax* :
    a. *tnow*_*h*_ = *tnext*_*h*_
    b. 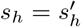
    c. Run code associated with the new state *s*_*h*_ and sample the next state transition and time (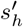, *tnext*_*h*_).
5. Set *t* = *tmax*.
6. Return to step 2 or quit.

When the agent is in a susceptible state and the accept-reject algorithm is being used to sample their time to infection, it is crucial that if the putative time of infection is rejected, the next time is set as *tnext*_*h*_ = *tmax* so that on the next time step, a new putative infection time is drawn. Because each agent updates themselves in continuous time within the while loop on step 4, the distribution of sampled times for internal events will be exact. Additionally, because each agent stores their own next state and time, so there is no need for a complex global event queue.

How in particular each agent samples from the tuple (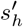, *tnext*_*h*_) in step 4c is left unspecified. Any exact sampling method for competing hazards is valid, and choice of method will depend on the problem at hand. Additionally we did not specify how to record model output; the method which provides the most granular output, and enabling survival analysis of results to verify the model, would delegate writing output to code associated with the event which causes the transition to each new state *s*_*h*_, in step 4c. If coarse-grained output is deemed sufficient, a natural place to track output is step 2, which already requires a for loop over all agents.

Fig 3 shows how the simulation algorithm samples from an SIR epidemic [25] with three agents over two time steps. The state space for each agent is *S* = {*S, I, R*}. At time *t* = (*n* − 1)Δ*t*, there are two infectious individuals and one susceptible individual. For susceptible individual 3, we additionally show their FOI *λ*_3_(*t*) in blue. Note that while agents may experience state changes at any point in time (agent 1 recovers during the first time step and agent 2 during the second time step), the FOI is only allowed to change at each time step when agents exchange information (shown by red arrows). During the first time step, agent 3 samples a putative time to infection which exceeds time *n*Δ*t*, so it is rejected. Agent 1 recovers sometime before the end of the time step, but is not allowed to update agent 3 until the start of the next step. When the next time step begins, after updating its FOI, agent 3 again samples a putative infection time, which occurs during that time step 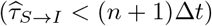, becoming infectious at that time. Additionally, agent 2 recovers sometime before the end of the time step.

**Fig 3.**
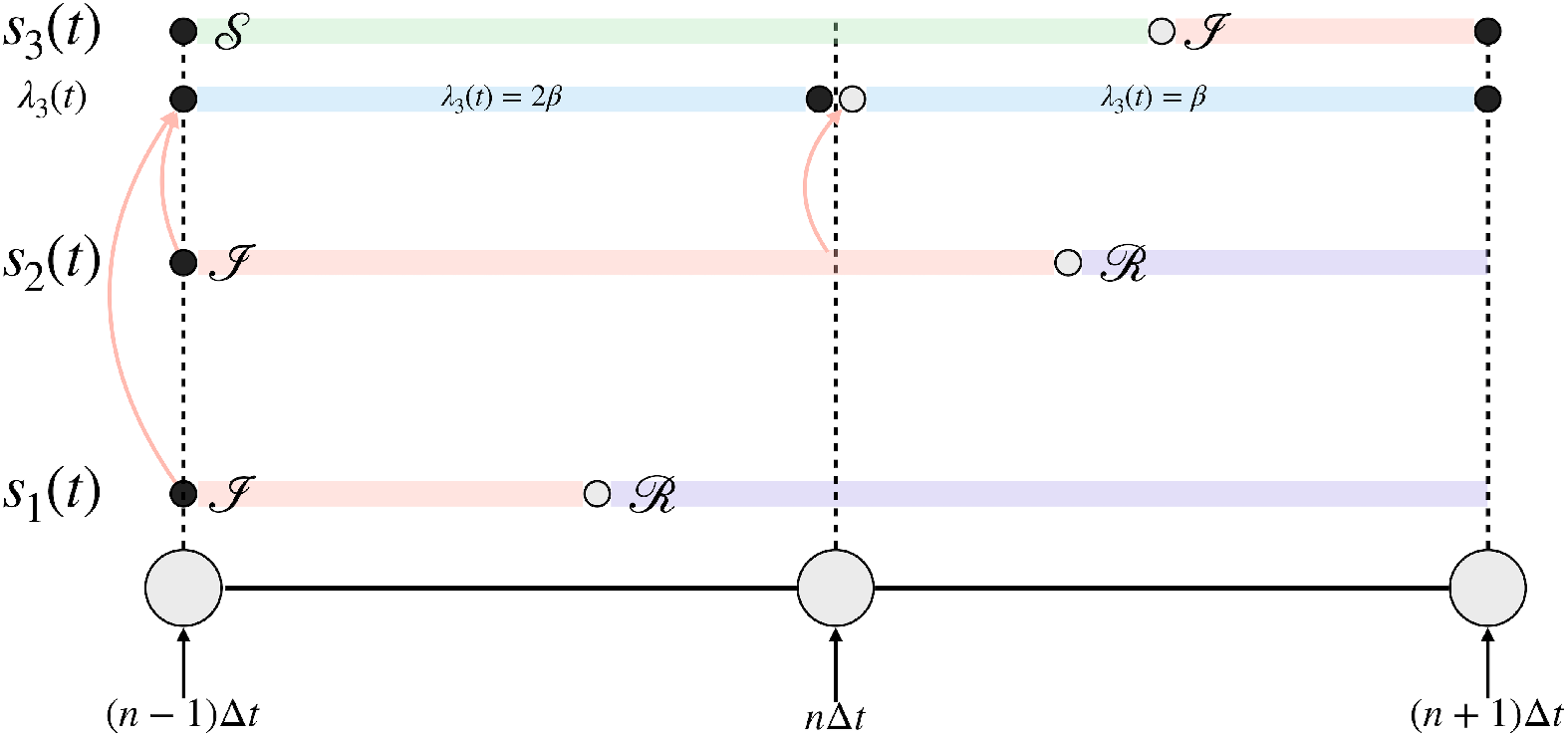
Graphical representation of simulation algorithm with three agents. Rows *s*_1_(*t*), *s*_2_(*t*), *s*_3_(*t*) are piecewise constant functions that give each agent’s state in the state space (*S, I, R*) at time *t*; each agents’ trajectory will be a piecewise constant function through state space. *S* (susceptible) is green, *I* (infectious) is blue, and *R* (recovered) is violet. The blue stripe *λ*_3_(*t*) gives the FOI on agent 3 (the only one to begin as susceptible).

While the diagram only shows a trajectory that overestimates the true FOI over each interval, if there was another *S* individual who transitioned to *I* during the time step their contribution would be left out and it would be an underestimate.

## Results

To illustrate use of our agent-based simulation method, we simulate a Markovian and non-Markovian SIR model. For both models we compare the sampled transition probability distribution from the ABM to that sampled from an exact stochastic simulation algorithm (SSA). We also compare samples from the ABM to closed form results on final epidemic size distributions, which provides exact analytic checks of algorithm accuracy. For the Markovian SIR model we can additionally compare the agent-based simulation to numerical solutions of the Kolmogorov forward equation (master equation) of the system, which gives the exact transition probabilities of the stochastic model. Because both these results provide a complete description of the probabilistic behavior of the stochastic models, they are more useful than comparing sampled trajectories (time series). However, we show some simple comparisons of sampled trajectories between exact stochastic simulation and the agent-based model in S3 Fig.

All of our code, written in R and C++ to reproduce all findings and figures in this paper is available at https://github.com/dd-harp/euler-abm.

### Markovian SIR Model

In Eq (1) we present the Kolmogorov forward equations (KFE) for the Markovian SIR model, following [25]. Because much recent research into methods for solving KFEs originates in statistical physics, which, unlike probability theory or mathematical epidemiology commonly uses *step operators*, we present the equations using step operators and work out a more familiar form, as presented in [26]. A brief introduction to using step operators to simplify writing KFEs for stochastic jump processes is available in S1 Appendix. The term **P**(*S, I, R, t*) is the probability for the system to be in state (*S, I, R*) at time *t*, such that *S* + *I* + *R* = *N*. We consider a simple mass action force of infection term to simplify the mathematics, so that the deterministic 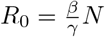, but the method is not restricted to simple mass action.

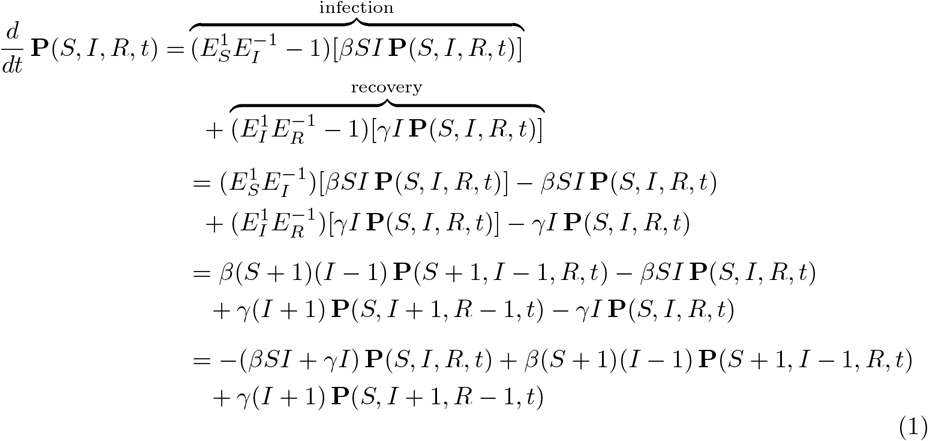

While solution of the KFEs via matrix exponentiation or, for small state spaces, direct numerical integration of the ODEs is possible, in order to evaluate the probability transition matrix over all possible 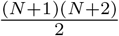 unique model states we use a recently developed technique based on continued fraction expansion which can calculate the state transition probability matrix directly [27, 28]. For an up to date review of methods for solving transition probabilities for common epidemic models, except the method of [28], see [29].

Solutions of Eq (1), computed via the continued fractions method are the “ground truth” against which we want to evaluate samples from our ABM method. For comparison to an exact SSA, we choose to implement the Modified Next Reaction Method (MNRM) of [30] to sample exact trajectories from the KFE because it is simple to code and easily extends to the non-Markovian case.

### Transition Probabilities

Solutions to the KFEs for the Markovian SIR model will give the exact transition probability matrix at a future time *t* given an initial state. That is, given some number of susceptible, infected, and recovered individuals at time 0, *S*(0), *I*(0), *R*(0), the KFEs can be solved to give the probability distribution over all possible states of the system at future time *t* ≥ 0, **P**(*S, I, R, t* | *S*(0), *I*(0), *R*(0)). This conditional probability distribution describes the exact probability law of the stochastic process; we use the method of [27], implemented in the R package, MultiBD [31] to solve for that distribution. We compare the exact distribution from solving Eq (1) to Monte Carlo simulation from the MNRM and ABM.

To assess the ability of the agent-based model to sample from the correct probability distribution over future states when simulating trajectories, we initiated a simulation with initial conditions *S*(0) = 60, *I*(0) = 10, *R*(0) = 0, *γ* = 1/3.5, and *R*_0_ = 2.5. We sampled 10^5^ trajectories from the ABM, exiting the simulation when the next event time would exceed *t* = 5, and using a time step Δ*t* = 0.01. We drew the same number of trajectories from the MNRM simulation algorithm so we could have a sense of how an exact stochastic sampler would approximate the true distribution. The results are visualized in Fig 4. We visualized the bivariate probability distribution over **P**(*S, I, t* = 5 | *S*(0), *I*(0)) using contours to represent curves of constant probability. Because of the constraint *N* = *S* + *I* + *R*, we do not lose any statistical information by disregarding *R*. Panel A compares the sampled MNRM (dashed contours) to the exact (solid contours), and Panel B compares the sampled ABM to the exact distribution. In both cases we observed very good equivalence between the sampled distributions and the exact distribution.

**Fig 4.**
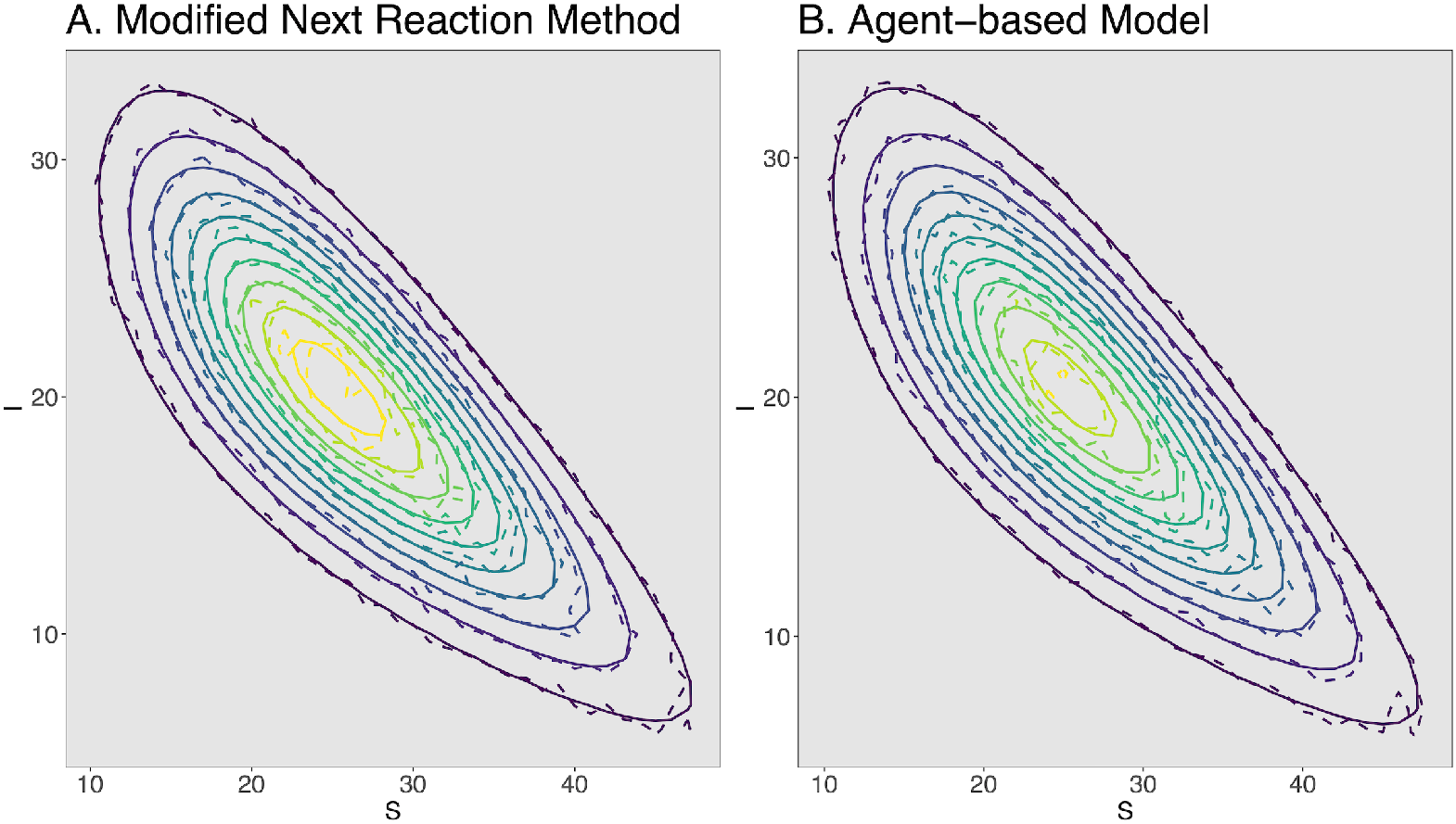
Comparison of exact transition probabilities to MNRM and ABM transition probabilities. Panel A: Comparison of MNRM (dashed contours) against exact probability distribution (solid contours), Panel B: Comparison of ABM (dashed contours) against exact probability distribution (solid contours). In both panels the x-axis and y-axis give the probability of having that number of susceptible and infectious individuals at *t* = 5, respectively.

### Effect of Time Step on Accuracy

We expect that as Δ*t* increases, the accuracy of the approximate ABM will deteriorate. We therefore evaluated the distribution **P**(*S, I, t* = 5 | *S*(0), *I*(0)) using the same parameters as the previous section, for a grid of time step sizes of 0.001, 0.005, 0.01, 0.025, 0.05, 0.075, 0.1, 0.5, 1. We chose this grid to span over several orders of magnitude, from extremely small values where we expect the ABM will be essentially exact, to a time step of one day, which for this specific setting is expected to be highly inaccurate. For each value of Δ*t*, we drew 2 × 10^5^ samples from the ABM to generate a Monte Carlo estimate of the transition probabilities. To calculate the exact transition probabilities, we used [31] to solve Eq (1).

For each time Δ*t*, we calculated absolute error as the sum of differences between the Monte Carlo estimate of the transition matrix and the exact transition matrix. Interestingly, although moving from the smallest step size of 0.001 to a more modest size of 0.1 spanned two orders of magnitude, absolute error changed little, from 0.045 to 0.059. Increasing Δ*t* to 0.5 and 1, however, swiftly increased absolute error to intolerable levels. We show a visual comparison of the effect of various step sizes on absolute error in Fig 5, where we plot the absolute difference between between each element in the bivariate distribution as a heatmap, visually revealing the method is highly accurate until large time steps of 0.5 and 1.

**Fig 5.**
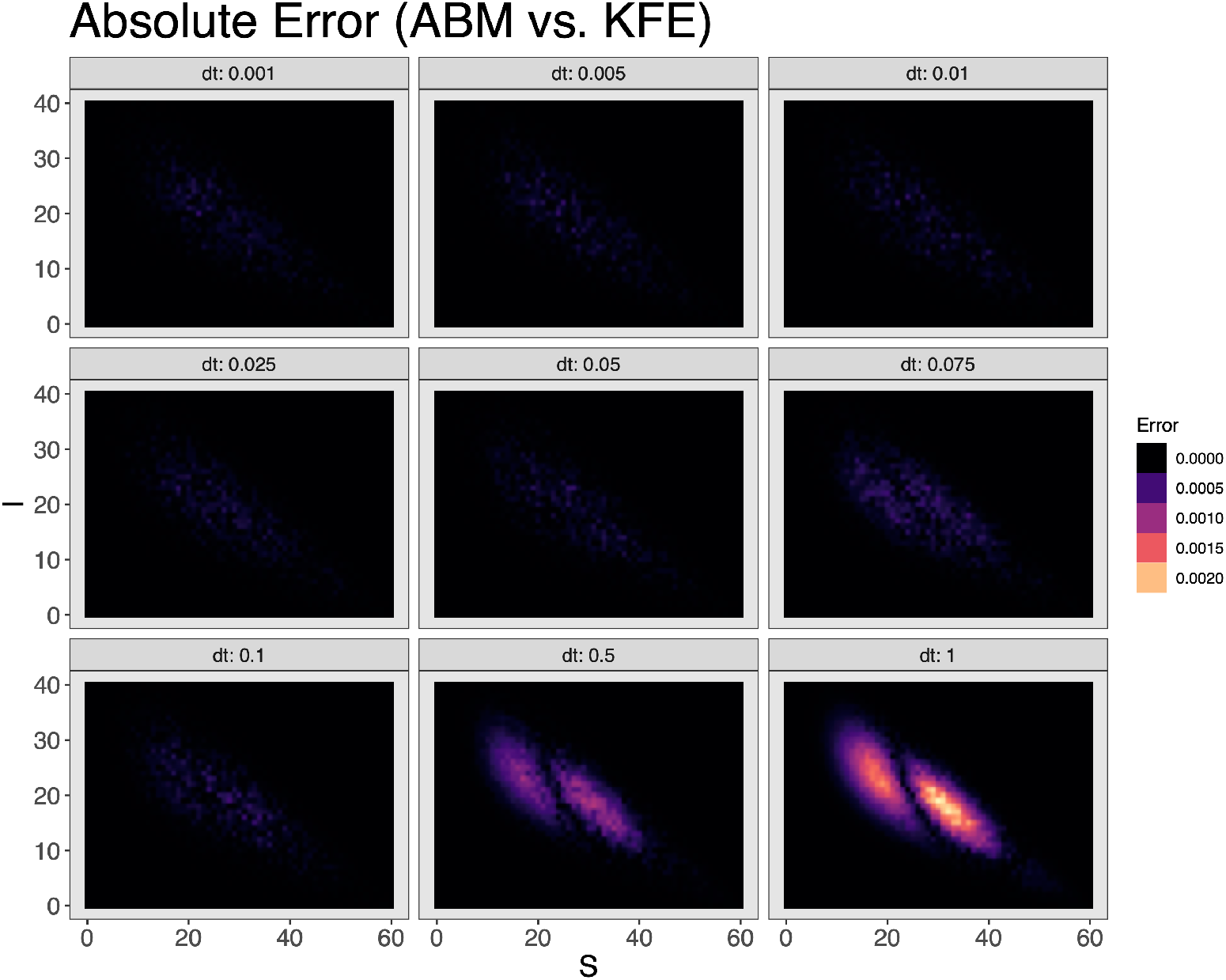
Absolute error between ABM and exact transition probabilities for different sized time steps. Panels show absolute error between transition probabilities calculated from the ABM versus exact distribution, from smallest time step (0.001) in the upper left to largest (1.0) in lower right. Darker areas correspond to small error while lighter regions correspond to higher error.

### Final Epidemic Size Distribution

An alternative to computing transition probabilities from Eq (1) for checking that our ABM is sampling from the correct process is to compare the final epidemic size distribution computed by Monte Carlo simulation to an exact analytic result. For SIR models, a closed form final epidemic size distribution was developed in [32, 33] and recently reviewed in [34]. This closed form solution for the distribution of final epidemic sizes is particularly valuable because it will be affected by the distribution of duration of infectiousness, meaning it provides a complete check on the the ability of a sampling algorithm to simulate the SIR model.

If the duration of infectious period *F* has a moment generating function (MGF) given by Ψ_*F*_ (*t*) = 𝔼[*e*^*tF*^] and where the initial states are given as *S*(0) = *N, I*(0) = *m*, the rate of effective contact by *λ*, then the final epidemic size vector is 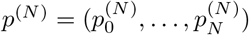 where 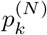 is the probability that *k* of the initial *N* susceptible individuals are infected when the epidemic ends. The probability 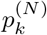 for any element of the vector is given in Eq (2) and a description of the solution is shown in S2 Appendix.

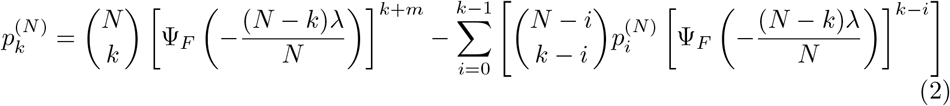

Because the equations use the MGF of the infectious period distribution, Eq (2) can compute final epidemic size distributions for both the Markovian and non-Markovian SIR model.

We compared the final epidemic size distribution sampled from the exact simulation algorithm (MNRM) and the ABM with Δ*t* = 0.01 to the exact closed form probabilities calculated from Eq (2) in Fig 6. We used initial conditions of *N* = 50 and *m* = 1, and sampled 10^4^ epidemic sizes from each stochastic simulation. The effective contact rate was calculated to give *R*_0_ = 2.5, and *γ* = 1/5. The exact probabilities are given by red dots, and the empirical probabilities are given by the purple horizontal lines; pointwise 95% confidence intervals were computed for each empirical probability by Wilson’s score method and the coverage interval is given as the shaded rectangle around the empirical probability [35]. In both cases the empirical distribution from stochastic simulation is nearly identical to the analytic probabilities, with remaining deviations due to Monte Carlo finite sample error.

**Fig 6.**
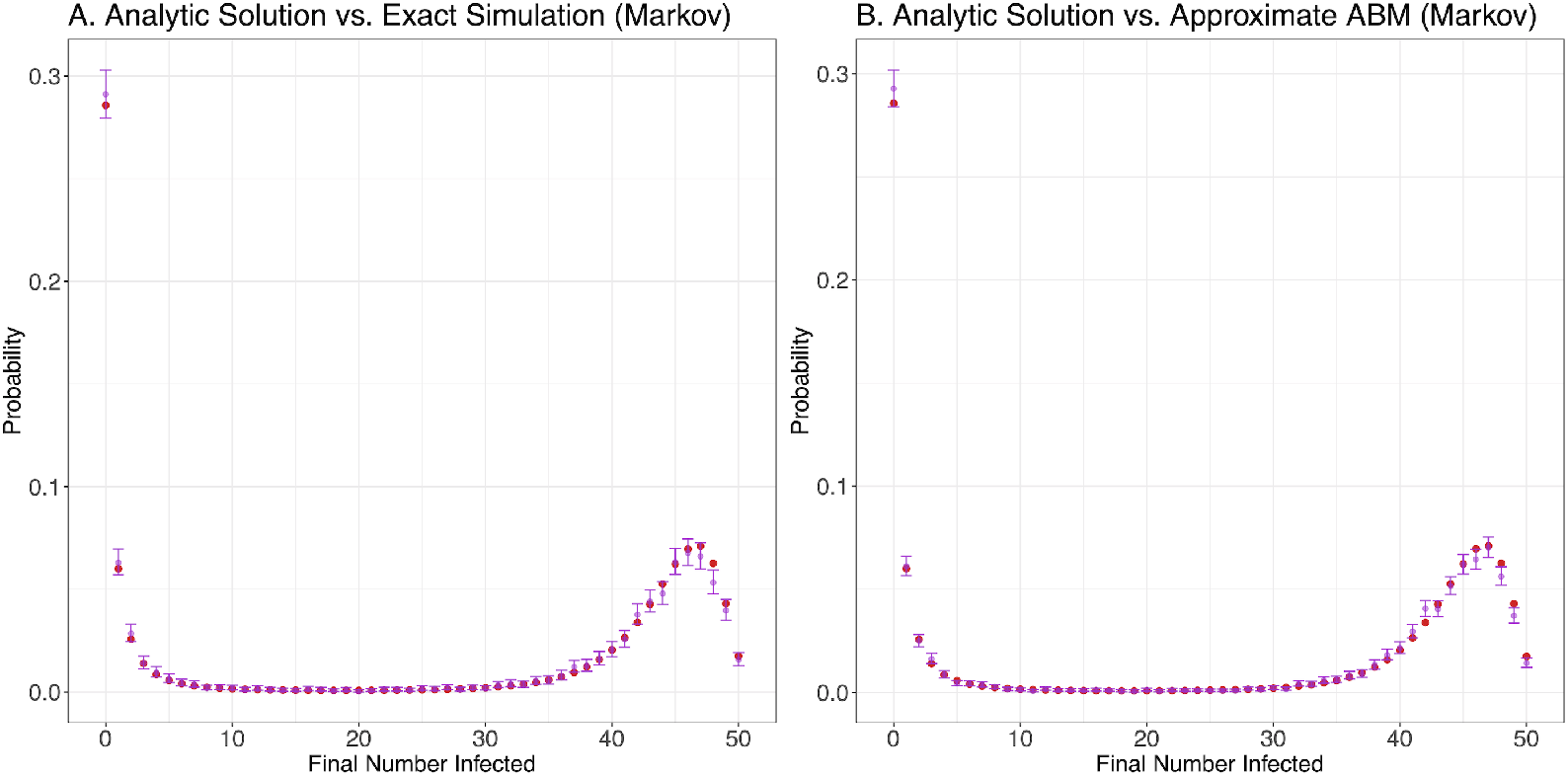
Final epidemic size distributions for Markovian SIR model. Panel A: Analytic final epidemic size distribution (red) versus empirical distribution (purple) from MNRM [30], Panel B: same, but empirical distribution (purple) from ABM. For each possible final size value we plotted the mean of simulation results as a purple dot with error bars indicating the pointwise 95% confidence interval from Wilson’s score method.

### Non-Markovian SIR Model

In Eq (3) we present the Kolmogorov forward equations for the non-Markovian (semi-Markov) SIR model, where the infectious period *τ* ∼*F* is a random variable that has a density function 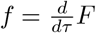, which may differ from an Exponential distribution. Because we assume that infection events still occur according to the points of a Poisson process, the contribution to the KFE from infection is the same as Eq (1).

However the contribution from the recovery term is more complicated. In particular, we need to introduce the two-time joint probability distribution **P**(*S, I, R, t*; *S*_*t*−*τ*_, *I*_*t*−*τ*_, *R*_*t*−*τ*_, *t* − *τ*), which is the joint probability of the system being in state (*S*_*t*−*τ*_, *I*_*t*−*τ*_, *R*_*t*−*τ*_) at a time *t* − *τ*, and in state (*S, I, R*) at a later time *t*. In Eq (3), the recovery term must sum over all possible states that the process could have been in *τ* units of time in the past, where *τ* ranges from [0, ∞), and is weighted by *f* (*τ*)*dτ*, the probability of an infectious period of duration *τ*. This means when a *S* particle becomes an *I* particle at time *t* − *τ* it *immediately* samples a time to recovery according to *f*. Those recovery events which complete at time *t* have probability *f* (*τ*)*dτ* of requiring that amount of time to do so.

In this sense, the recovery events in the system are still controlled by the points of the Poisson process generating infection events, but represent delayed effects of that event. An excellent description of stochastic systems with delayed effects is given in [36]; another slightly different approach to these systems is described by [37].

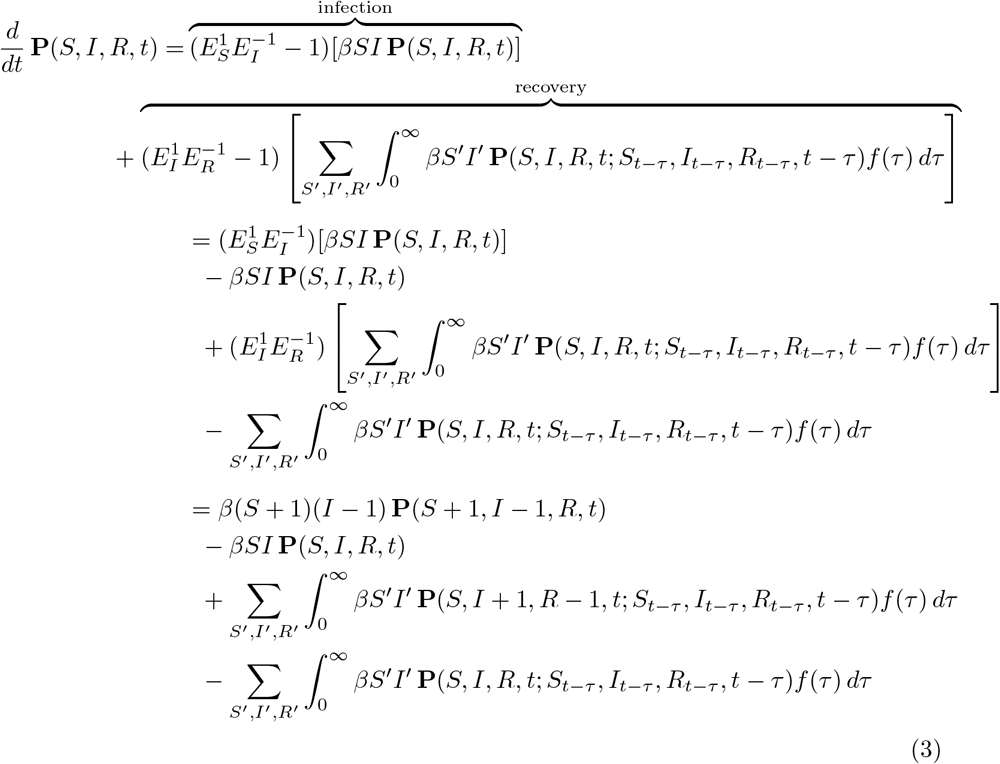

The final non-Markovian Kolmogorov forward equation has four terms. The first and second terms represent infection events that will push the state into, and out of (*S, I, R*), respectively. The third term represents infection events that fired some time *τ* in the past and whose delayed recovery event will push the process into state (*S, I, R*) at time *t*. Likewise the final term represents those infection events whose delayed recovery events are just about to push the process out of that state.

To draw exact samples from the non-Markovian SIR model, we used the MNRM from [30] for arbitrarily distributed delays. We modified Algorithm 7 such that the delayed reaction channels store completion events as a priority queue implemented as a binary heap, after drawing the delay, *τ* from the appropriate distribution.

### Transition Probabilities

To assess the ability of the agent-based model to sample from the correct transition probability distribution over future states when simulating trajectories from the non-Markovian model, we initiated a simulation with conditions *S*(0) = 60, *I*(0) = 10, *R*(0) = 0, *γ* = 1/3.5, and *R*_0_ = 2.5. We sampled 10^5^ trajectories from the ABM, exiting the simulation when the next event time would exceed *t* = 5, and using a time step Δ*t* = 0.01. We drew the same number of trajectories from the MNRM simulation algorithm in order to compare approximate transition probabilities from the ABM to an exact sampler. Unlike the Markovian SIR model, the non-Markovian KFEs in Eq (3) are analytically and numerically intractable, so the comparison in this section is only between the exact MNRM and ABM. The results are displayed in Figure 7. We visualized the bivariate probability distribution over **P**(*S, I, t* = 5 |*S*(0), *I*(0)) using contours to represent curves of constant probability. We observed a very good equivalence between the transition probabilities sampled via the approximate ABM (dashed contours) and exact sampler (solid contours).

**Fig 7.**
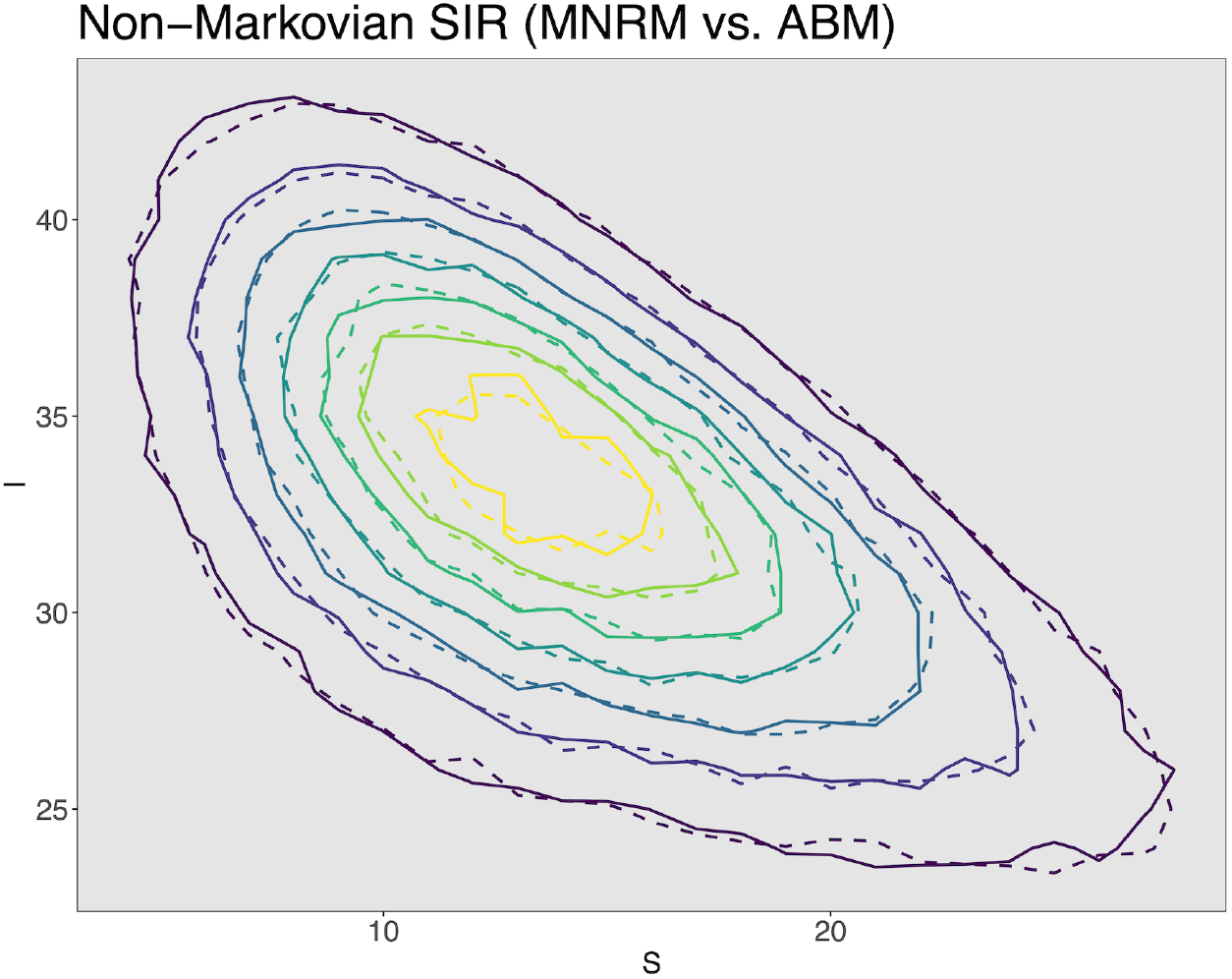
Comparison of exact transition probabilities to MNRM and ABM transition probabilities. Comparison of transition probability distribution sampled from MNRM (solid contours) to distribution sampled from the ABM (dashed contours). The x-axis and y-axis give the probability of having that number of susceptible and infectious individuals at *t* = 5, respectively.

### Final Epidemic Size Distributions

As a further verification that our approximate ABM correctly sampled trajectories for the non-Markovian model, we used Eq (2) to compute exact final epidemic size distributions for the non-Markovian SIR and compared both the MNRM samples and ABM samples to the closed form solution. The delay distribution (*f* (*τ*) in Equation 3) was a Gamma distribution, with a mean of 5 days and standard deviation of 0.5 days, giving a distribution peaked near the mean with most of the probability mass concentrated between 4 and 6 days. We chose a Gamma distribution because it has an analytically tractable moment generating function, so that the closed-form final epidemic size distributions are easy to compute.

We calibrated *β* to give *R*_0_ = 1.85, and sampled 10^4^ epidemic sizes from each simulation method. The ABM used a time step size Δ*t* = 0.01. As before, *N* = 50 and *m* = 1. The procedure used to construct pointwise confidence intervals was the same as used in the Markovian example. Fig 8 shows that our approximate ABM with the chosen time step is able to sample from the closed form probability distribution with high accuracy (Panel B), and its performance is indistinguishable from an exact sampler (Panel A).

**Fig 8.**
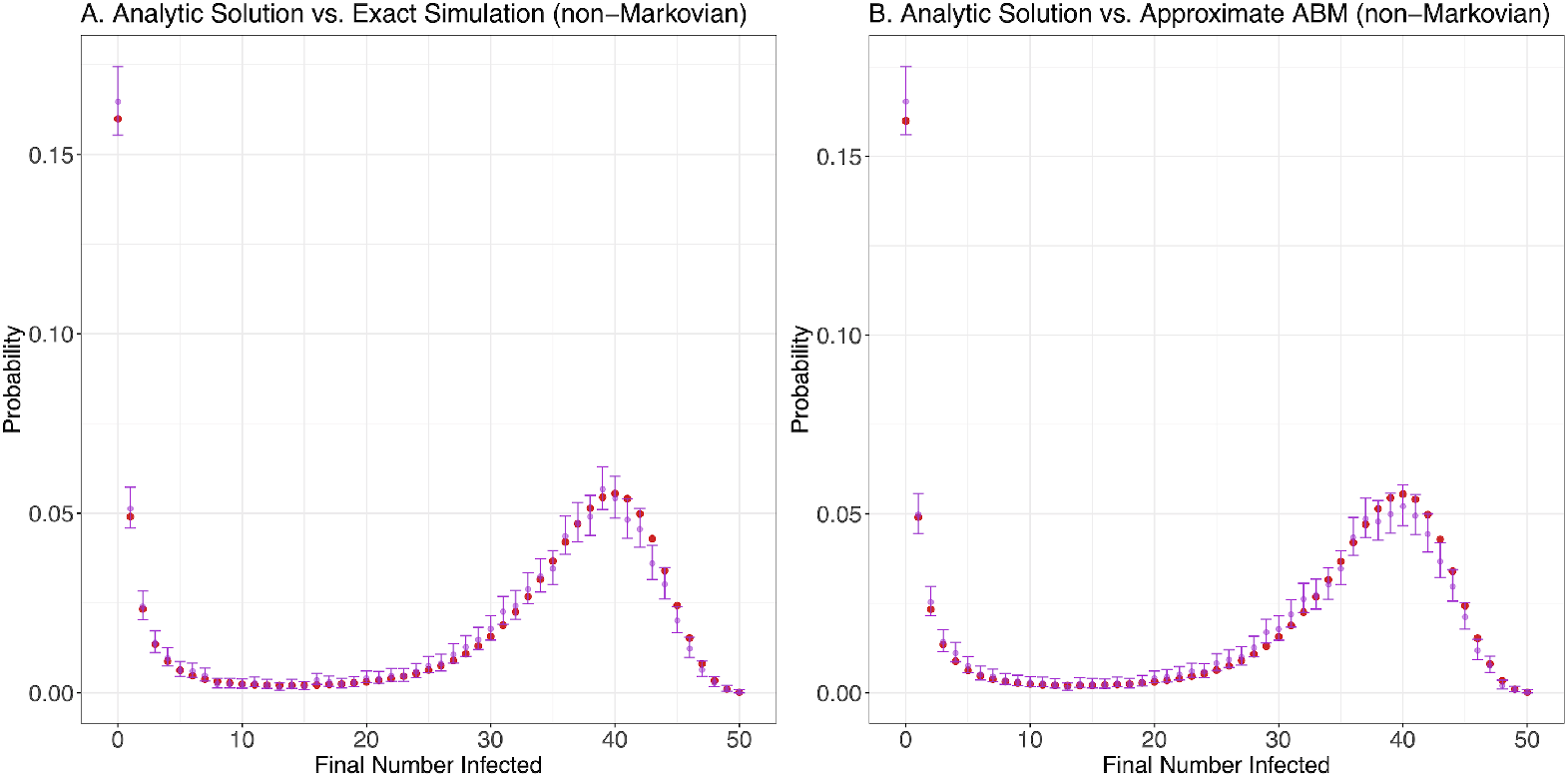
Final epidemic size distributions for non-Markovian SIR model. Panel A: Analytic final epidemic size distribution (red) versus empirical distribution (purple) from MNRM [30], Panel B: same, but empirical distribution (purple) from ABM. For each possible final size value we plotted the mean of simulation results as a purple dot with error bars indicating the pointwise 95% confidence interval from Wilson’s score method.

## Discussion

We have presented a practical approach to simulating ABMs with attractive properties. One interesting consequence of using a time step is that integration with other models becomes easier. If certain hazards depended upon, say a mosquito model with some discrete time step, that information could be exchanged between models at each step. As the model is formulated with continuous hazards, the size of a time step can vary over a simulation run, potentially widening applicability of the method. Furthermore, its generality means there are several interesting avenues for further development.

A clear next step is a method for bounding approximation error during a simulation run, which would allow for development of methods for adaptive selection of Δ*t*. While the ABM formulation means the model’s computational structure is different (disaggregated) compared to chemical kinetic simulations, hazard functions can could still be queried by looping over all individuals in the simulation. Because the discrete time step only approximates dependent events, agents without any enabled dependent events could be skipped. A first approach at adaptive selection of Δ*t* may look somewhat similar to the straightforward methods presented in [21], and there is a wealth of citing literature that could be used for further development.

Another interesting avenue for future research is investigating how to handle multiple types of interaction which occur on different time scales. Because in the example SIR models presented here, agents only interact through a single dependent event, which is assumed to occur at the points of a Poisson process, all interactions occur on the same time scale. However, if multiple types of dependent interaction were simulated (frequent contacts between close friends versus rare meetings between acquaintances, for example), total error induced by a choice of Δ*t* would depend on contributions from approximation error from all interaction types. While chemical kinetics has developed several methods for dealing with reactions occurring on very different characteristic time scales (see [38] for an example), it is not clear what is the best way to deal with separation of time scale in the ABM. A naive approach would involve slower interaction terms being updated only at some integer multiple of Δ*t*, but further work should be undertaken to characterize the best way to choose such an updating scheme suitable for multiple time-scales, with the caveat that appropriate schemes would be highly problem dependent.

Our method is not restricted to pure jump stochastic processes. One generalization of a pure jump process is to assume that the system can be described by coupling continuous state variables to the discrete variables such that between jump times, the continuous part evolves according to a differential equation. Such systems are known as piecewise deterministic Markov processes (PDMP) when the distribution of inter-event times is given by an nonhomogeneous Poisson process whose intensity may depend on the continuous variables [39]. This model representation may be natural to simulate multi-scale models, where within-host immune dynamics follow a differential equation model, like the method proposed in [40] to model host-pathogen interaction. In the case where the deterministic dynamics are fully internal to each agent the implementation simply involves solving a differential equation for each agent to sample their next state and time. More complex internal dynamics, such as allowing the continuous state to follow a stochastic differential equation, can be implemented in the same way, as long as the next jump time for each agent’s discrete state is computable from the solution of the process (for example, from a first hitting time). These models can be useful to integrate results from medical survival analysis [41].

The computational methods for epidemiological problems have borrowed heavily from other computational disciplines, but some of the unique features of epidemiological systems would benefit innovation and algorithms to addressing the kinds of problems that arise. In particular, for simulation of complex epidemiological models, agent-based models are a useful alternative to other modeling techniques, and can be both flexible and extensible. In this work we have presented a generic algorithm to sample trajectories from agent based models, for agents which may experience dependent or internal events, parameterized by hazard functions. Approximation of a subset of event hazards, those that depend on multiple agents’ state, speeds up simulation. A key contribution of our ABM simulation technique is that it will converge to a continuous-time stochastic jump process in the limit of small step sizes. Our intention is not to argue that models without a limiting interpretation are wrong, but that in many practical cases being able to connect the ABM to a well defined stochastic process is valuable, such as for parameterization from results of survival analysis or for verification of simulation software. Construction from continuous-time hazards means that ABMs developed in this framework can be more clearly linked to other results in stochastic simulation, easing inclusion of various elaborations relevant to epidemiological simulation, such as time-varying hazards [42, 43]. We hope that the simplicity of the method presented in this paper can help researchers respond more quickly to urgent situations where stochastic models are required.

## Supporting information

**S1 Appendix. Brief Description of Step Operators**. Gives a brief introduction to the use of step operators when writing the Kolmogorov forward equations for stochastic jump processes.

**S2 Appendix. Final Epidemic Size Distributions**. Gives more details on the closed form probability for final epidemic size distribution with arbitrary distribution of infectious period.

**S1 Fig. Truncated Exponential distribution of infection time on time step where the** 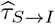 **is accepted**. An Exponential distribution truncated at Δ*t* with rate parameter *λ* has the density 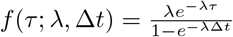.

**S2 Fig. Nonhomogeneous diurnal intensity function**. Continuous intensity function 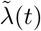 (red solid line) and piecewise approximation *λ*(*t*) (purple step function; approximation used for simulation is on much finer time-step, coarse approximation is purely for visual effect).

**S3 Fig. Comparison of simulation trajectories from exact stochastic simulation and ABM**. Panel A: Markovian SIR trajectories. Panel B: non-Markovian SIR trajectories. For each panel we drew 10^4^ trajectories from the MNRM and ABM simulation algorithms and summarized the results by plotting the mean and 95% simulation interval. The MNRM trajectory is denoted by dashed lines and the ABM by solid lines.

## Acknowledgments

We thank Robert C. Spear for discussions on early drafts of this work.

## Brief Description of Step Operators

Step operators are a convenient way to simplify the expression of Kolmogorov forward equations (master equations; henceforth KFE) for jump processes. We introduce them here in the context of continuous-time Markov chains, following the exposition of Toral and Colet 2014. Mathematically, the step operator *E* is a linear operator on a function *f* (*n*) where *n* ∈ ℤ. Because the probability flux on the right hand side of the differential KFE has contributions from all states with nonzero transition probabilities, and states are expressed as vectors of integers, the step operator *E* can be used to write KFEs. The main advantage of using step operators to write the KFE is that one no longer needs to keep track of what specific elements of the probability distribution the flux is flowing in and out of, one simply needs to keep note of how many “particles” of each type (corresponding to specific positions in the state vector) are being created and destroyed for each event that is allowed to change system state.

The step operator *E* is defined as below; for events which may cause the creation or destruction of more than one particle, note that *E* is defined for powers *l*. Most important for actually writing KFEs is the final relation, (*E*^*l*^ − 1), which simplifies the inbound and outbound probability fluxes into a single term.

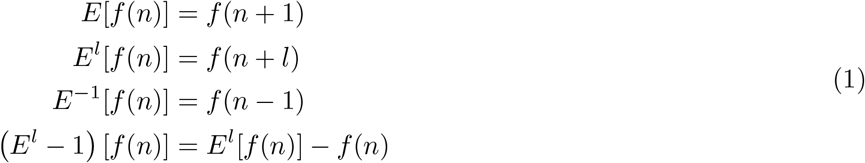

When writing KFEs with the step operator defined in Equation 1, note that to represent an event which creates *l* particles, one would use (*E*^−*l*^ − 1) [*f* (*n*)]. This is because the probability flux will include an incoming component from the state with *l* fewer particles, and the outbound flux from the current state.

Finally, for systems with more than one type of particle (the state space is a vector with more than one element), we can compose step operators to represent events which simultaneously change multiple particle counts simultaneously. Examples include predation in the stochastic Lotka-Volterra model, where predation simultaneously increases the count of predators and decreases the count of prey, or infection in simple epidemic models, which decrease the count of susceptibles and increases the count of infecteds. For an event which changes the count of type 1 and 2 particles by *l*_1_ and *l*_2_, one would use Equation 2. To evaluate 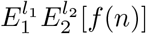, one applies the operators on *f* (*n*) one after the other.

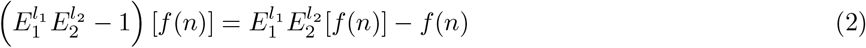

To illustrate use of the step operator *E*, we demonstrate their use on a Lotka-Volterra model. In this model there are two types of particles, prey *X* and predators *Y*. There are three events, which can be written using notation from chemical kinetics as follows:

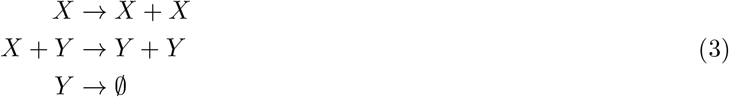

These correspond to reproduction of prey, predation, and death of predators. At any point in time, the probability of the state *n*_1_ prey (*X*) and *n*_2_ predators (*Y*) at time *t* is denoted as *P* (*n*_1_, *n*_2_; *t*). Reproduction of prey has per-capita rate *β*. Predation occurs according to simple mass-action with reaction constant *γ*. Finally, death of predators occurs with per-capita rate *µ*.

Now we write out the KFE and repeatedly apply the step operator until we get back to the familiar component-wise form.

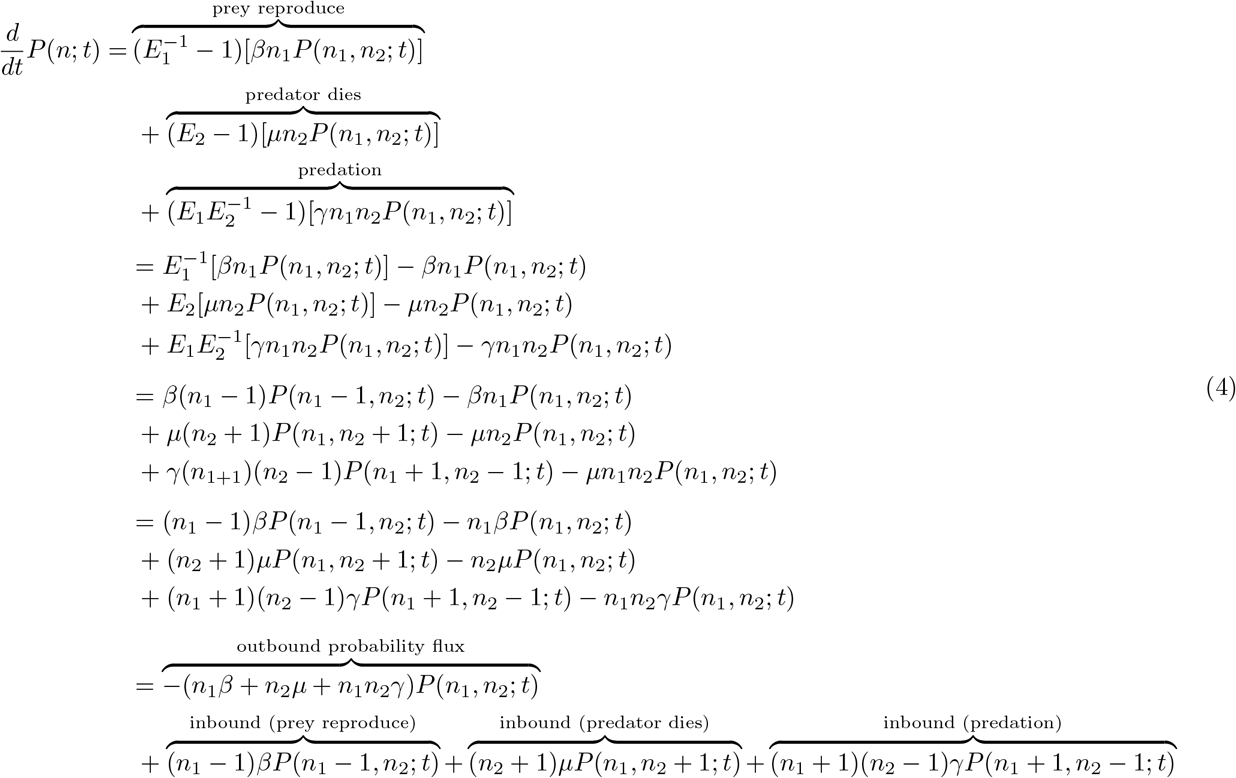

## Supplemental Information: Derivation of Final Epidemic Size Distributions

Another metric to check that our ABM is sampling from the correct process is to compare the final epidemic size distribution computed by Monte Carlo simulation of the ABM to an exact analytic result. We rely on the method presented in (Sellke 1983) and well described in (Andersson and Britton 2012) to compute final size distributions for SIR epidemic models with general distributions of infectious periods.

For a SIR epidemic where infectious periods *F* have a moment generating function (MGF) which is Ψ_*F*_ (*t*) = 𝔼[*e*^*tF*^] and where the initial states are given as *S*(0) = *N, I*(0) = *m*, the rate of effective contact by *λ*, then the final epidemic size vector is 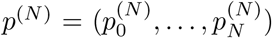 where 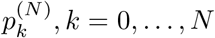 is the probability that *k* of the initial *N* susceptibles are infected when the epidemic ends. From (Ball 1986), these probabilities are related by the recursive equation:

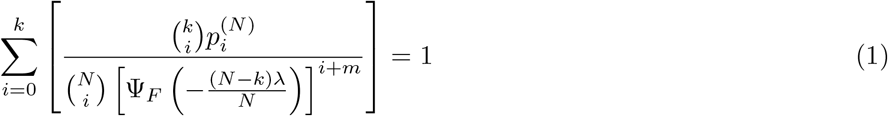

Using Equation 1, we calculate an explicit recursive formula for the probabilities 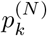, *k* = 0, …, *N*. We begin by using 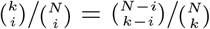 to get:

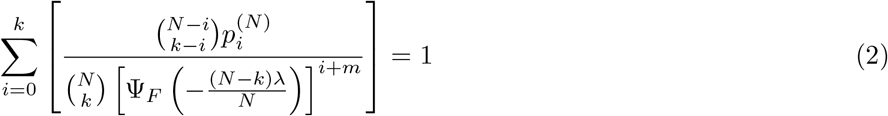

Then pull 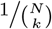 out of the sum and multiply both sides by 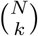:

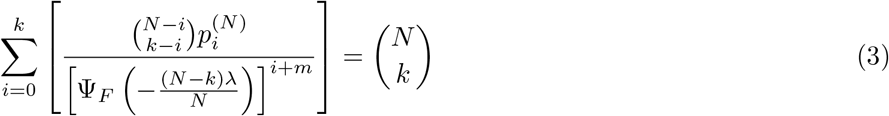

Next pull out the final summation of the left hand side:

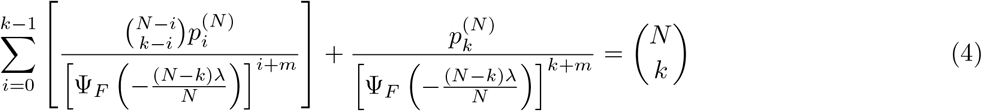

Now multiply all terms through by 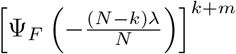, noting that we can move the power in the denominator up to the numerator by adjusting the power by (*k* + *m*) − (*i* + *m*) = *k* − *i*:

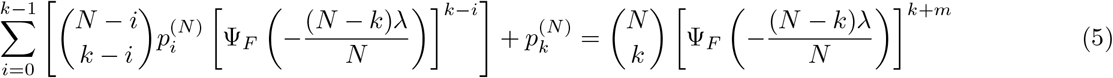

Finally, solve for 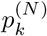, the desired probability:

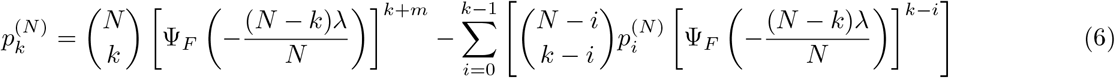

In order to turn Equation 6 into an algorithm, we note that when *k* = 0 the sum drops out and the equation simplifies to:

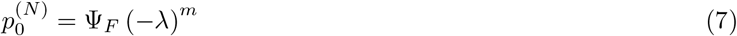

Probabilities for *k* = 1, …, *N* can then be solved for recursively. It should be noted that for *N* much larger than 100, the algorithm given by Equation 6 suffers from numerical instability, and so should only be used for small populations. Because the equations are defined solely in terms of the MGF of the infectious period distribution, the probabilities from Equation 6 can be used for both the Markovian and non-Markovian SIR model.

## Supplemental Information: Figures

**Figure 1:**
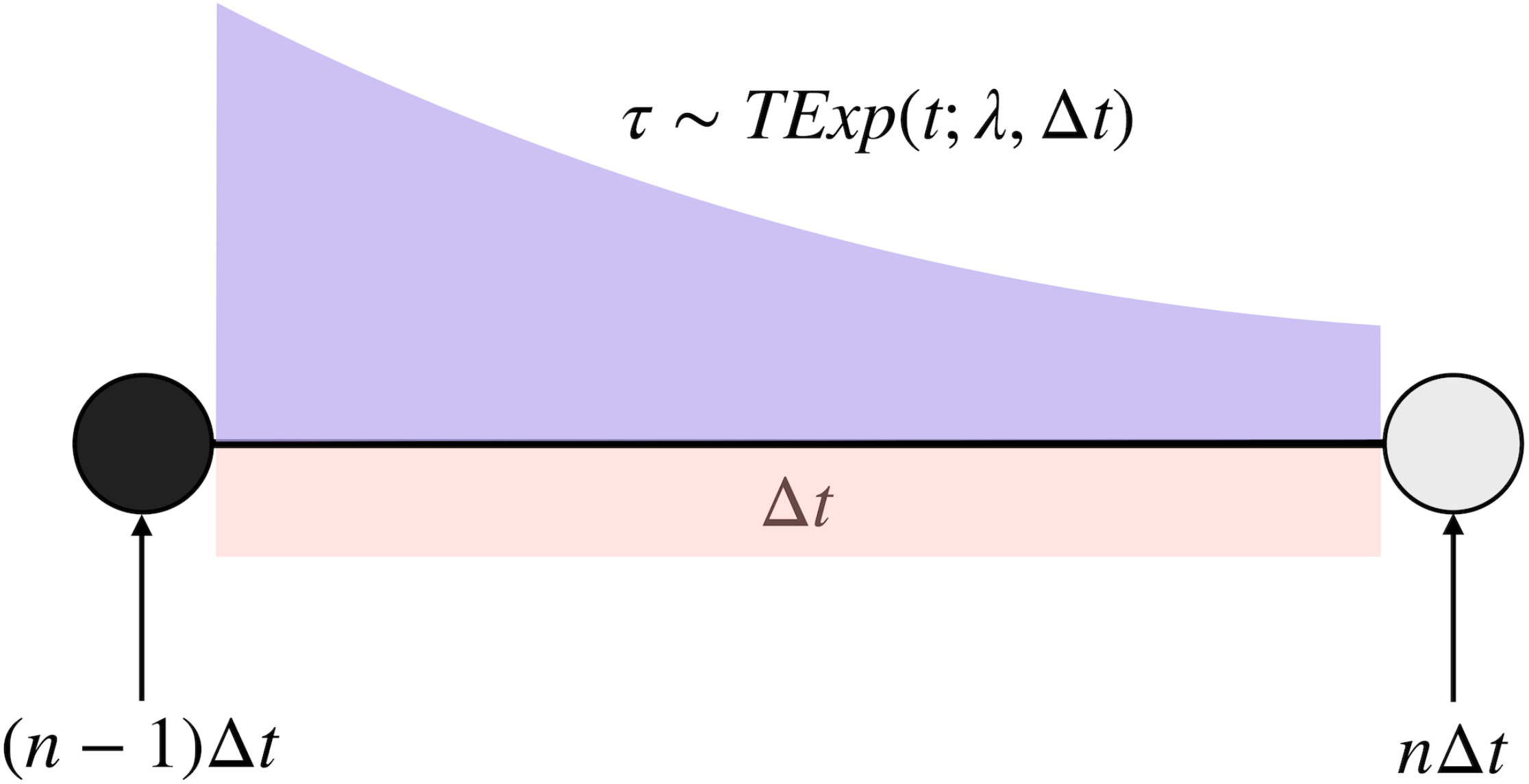
**Truncated Exponential distribution of infection time on time step where the 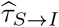 is accepted.** An Exponential distribution truncated at Δ*t* with rate parameter *λ* has the density 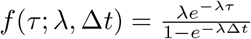.

**Figure 2:**
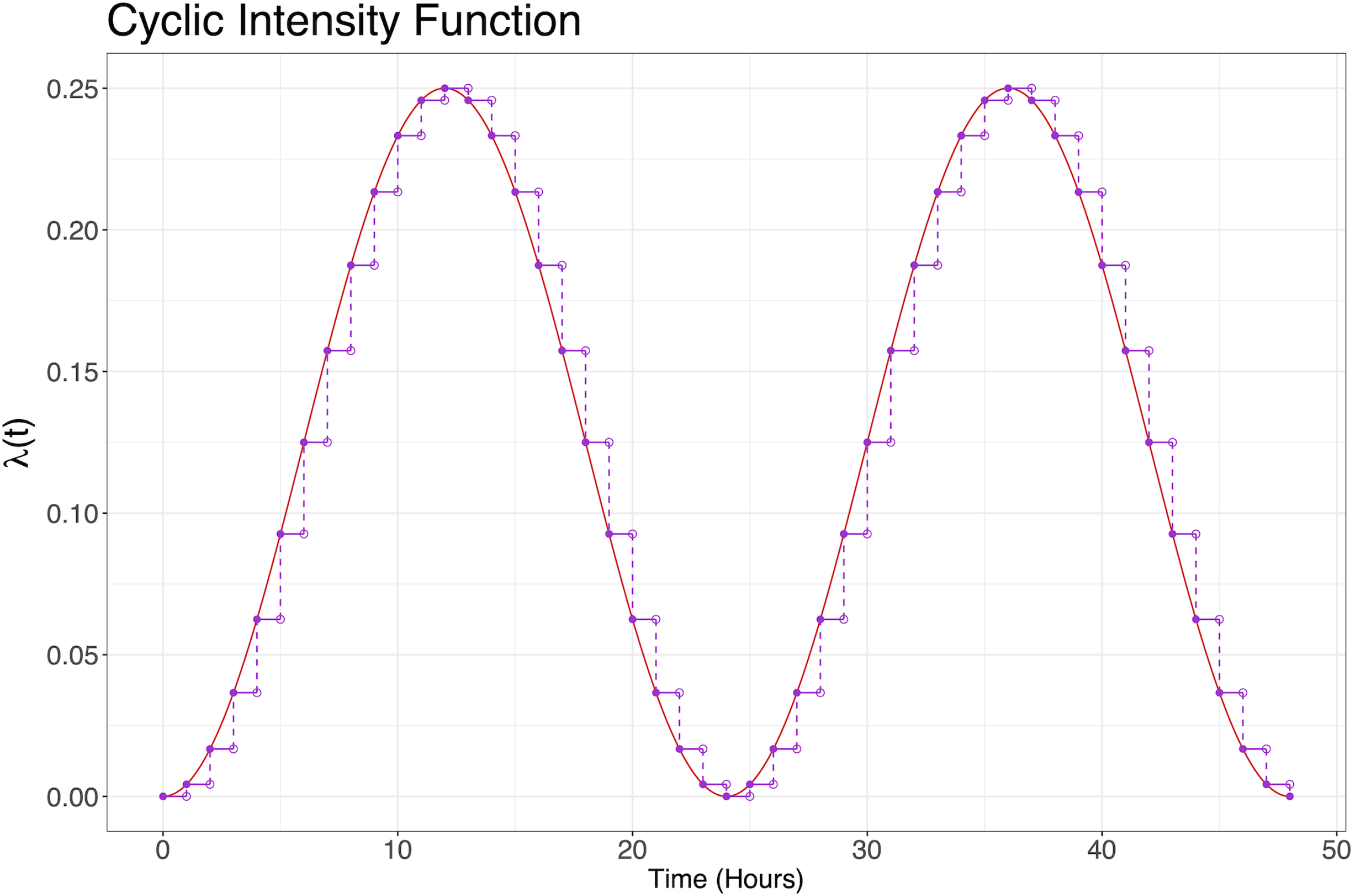
Nonhomogeneous diurnal intensity function. Continuous intensity function 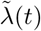 (red solid line) and piecewise approximation *λ*(*t*) (purple step function; approximation used for simulation is on much finer time-step, coarse approximation is purely for visual effect).

**Figure 3:**
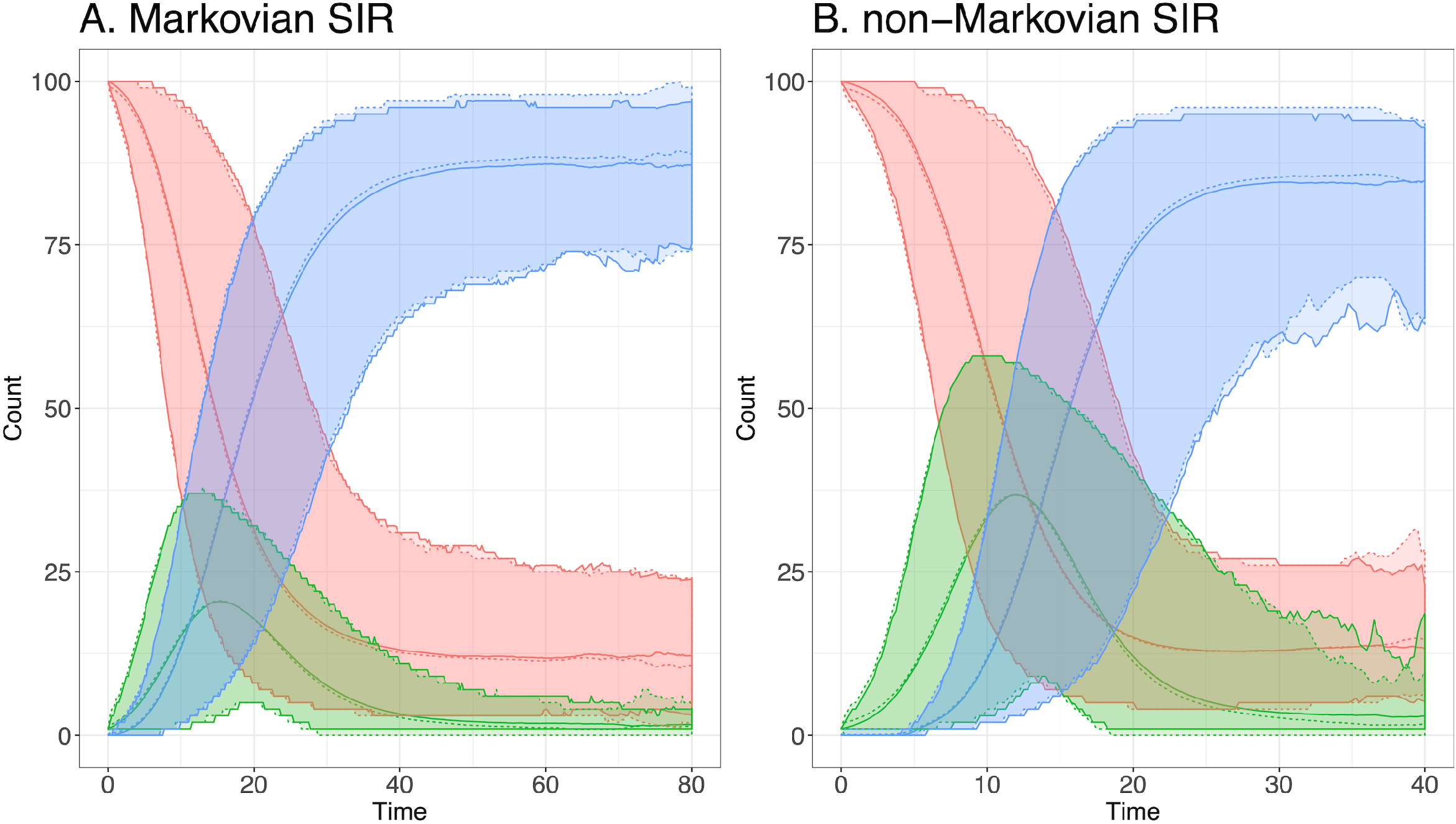
Comparison of simulation trajectories from exact stochastic simulation and ABM. Panel A: Markovian SIR trajectories. Panel B: non-Markovian SIR trajectories. For each panel we drew 10^4^ trajectories from the MNRM and ABM simulation algorithms and summarized the results by plotting the mean and 95% simulation interval. The MNRM trajectory is denoted by dashed lines and the ABM by solid lines.

